# Neural structure of a sensory decoder for motor control

**DOI:** 10.1101/2020.10.22.350371

**Authors:** Seth W. Egger, Stephen G. Lisberger

**Affiliations:** Department of Neurobiology, Duke University School of Medicine, North Carolina 27710, USA

## Abstract

We seek to understand the neural mechanisms that perform sensory decoding for motor behavior, advancing the field by designing decoders based on neural circuits. A simple experiment produced a surprising result that shapes our approach. Changing the size of a target for smooth pursuit eye movements changes the relationship between the variance and mean of the evoked behavior in a way that contradicts the regime of “signal-dependent noise” and defies traditional decoding approaches. A theoretical analysis leads us to conclude that sensory decoding circuits for pursuit include multiple parallel pathways and multiple sources of variation. Behavioral and neural responses with biomimetic statistics emerge from a biologically-motivated circuit model with noise in the pathway that is dedicated to flexibly adjusting the strength of visual-motor transmission. Flexible adjustment of transmission strength applies much more broadly to issues in sensory-motor control such as Bayesian integration and control strategies to optimize motor behavior.

## Introduction

The complex circuits of the brain transform sensory inputs into appropriate motor outputs. However, the brain is imperfect, and motor outputs vary from trial to trial even for identical sensory inputs. The variation has a distinct form, leading to well known psychophysical “laws”. Fitts law describes a proportional increase in movement variation with the speed or size of the movement^1^. The Weber-Fechner law states that sensory discrimination thresholds grow in proportion with stimulus amplitude^2^. Both laws are classically attributed to “signal dependent noise”, where noise in the brain increases with the amplitude of sensory or motor signals.

Signal-dependent noise has a long history in models of sensory-motor behaviors. To capture the psychophysical laws, models of sensory-motor transformations often take a “black box” approach, where application of a decoding equation to sensory representations generates motor outputs^3–8^. Signal-dependent noise is built into behavioral output either through the properties of the sensory representation^9, 10^ or noise in the motor command^11, 12^. It is assumed, therefore, that noise is absent from the varied and complex decoding circuits that intervene between sensory representations and motor coordination.

In contrast to the view that decoding circuits are free of noise, the responses of neurons making up the circuits between sensory and motor systems vary considerably from trial to trial^13^. The likely presence of decoder noise challenges the “black box” design of decoding models, as the non-sensory, non-motor neurons that perform the computations required of decoding are proposed to contribute to behavioral variation^14, 15^. Further, the ability of models that delegate noise exclusively to sensory or motor sources to explain behavioral variance becomes more limited as behavioral tasks become more complicated^16–18^. Our goal is to understand the neural computations that generate sensory-motor behavior, and doing so requires opening the “black box” to consider the computations performed by specific neural pathways as they become engaged during increasingly complex behaviors. The key question that arises from this approach is not whether the responses of neurons in the decoder vary (they do), but rather if the variation that arises in each neural pathway contributes uniquely to behavioral variation as a consequence of the pathway’s specific function in sensory-motor transformations.

We approach a mechanistic understanding of sensory decoding by studying visually-guided smooth pursuit eye movements, a system where we know much about the underlying neurophysiology. The extrastriate middle temporal visual area (MT) provides sensory signals that drive the behavior^19–21^ and several strong empirical observations link the responses of MT neurons to behavioral variation. First, the limits on sensory precision by MT neurons are similar to pursuit eye movements^22^. Second, the properties of MT neurons are appropriate to transform the fluctuations of individual neurons into signal-dependent noise^23^. Finally, trial-by-trial “neuron-behavior” correlations between the responses of individual MT neurons and the eye velocity in the initiation of pursuit argue that correlated sensory noise contributes to motor variation, and places limits on how much variation is added downstream^24^. The robust link between responses in MT and motor behavior has led to a canonical model where downstream circuits simply decode MT responses and the fluctuations in MT neurons are the primary driver of variation in pursuit.

We start by presenting the simple experimental observation that changing target size breaks the traditional psychophysical laws of signal-dependent noise. The observation was unexpected and cannot be explained by what we know about sensory or motor systems, motivating a consideration of signal and noise in the circuit pathways that make up the sensory decoder. Pursuit eye movements have a neural substrate with at least two pathways radiating from the sensory representation before converging onto the final motor circuits^25^. Anatomically, the output from area MT is transmitted both through fairly direct ponto-cerebellar pathways and through a cortico-cortical circuit that involves the smooth eye movement region of the frontal eye fields (FEF_sem_). FEF_sem_ exerts profound control over pursuit behavior by modulating the strength or “gain” of visual-motor transmission^26^. The “gain control” pathway seems to afford flexible, context-based visual-motor transformations as opposed to reflex-like input-output behavior^25, 27–29^. The combination of multiple anatomical, physiological, and behavioral observations leads directly to the proposition that an accurate decoding model will include a pathway for gain control that makes separate and independent contributions to both signal and variation.

We present a new model of the sensory-motor decoding that respects the neuroanatomy of pathways downstream from MT, the tight link between neural activity in MT and FEF_sem_^30^, and extensive neurophysiological and behavioral data^24, 31^. The decoder includes parallel pathways for (1) computing the gain of visual-motor transmission and (2) estimating target speed. Noise in the gain pathway not only accounts for the breakage of traditional psychophysical laws by changes in target size, but also captures competing constraints provided by physiological observations. The gain pathway is a critical piece of this specific neural system, and also applies much more broadly to issues central to optimized sensory-motor control. For example, Bayesian integration^29, 32, 33^ and control strategies to optimize motor behavior^12, 34, 35^ all depend critically on controlling the relative gains of different influences on behavior.

## Results

### The relationship between behavioral variance and mean changes with the size of the pursuit target

We start by presenting a simple experiment with results that contradict previous conceptions of the relationship between the variance and mean of motor behavior. We analyzed the eye movements of monkeys that were rewarded for smoothly pursuing patches of moving dots of different sizes. In each trial, a patch of dots centered on the fovea moved within a stationary aperture for 150 ms before moving along with the aperture across the screen. Target speeds were selected randomly from trial-to-trial from 4, 8, 12, 16, and 20 deg/s with equal probability (Figure 1A, top). Target sizes were selected randomly with equal probability from diameters of 2, 6, and 20 deg (Figure 1A, bottom). Target speed and size strongly influenced mean eye speeds over the first 250 ms after the onset of target motion (Figure 1B), as expected based on previous experimental results^36–38^.

**Figure 1.**
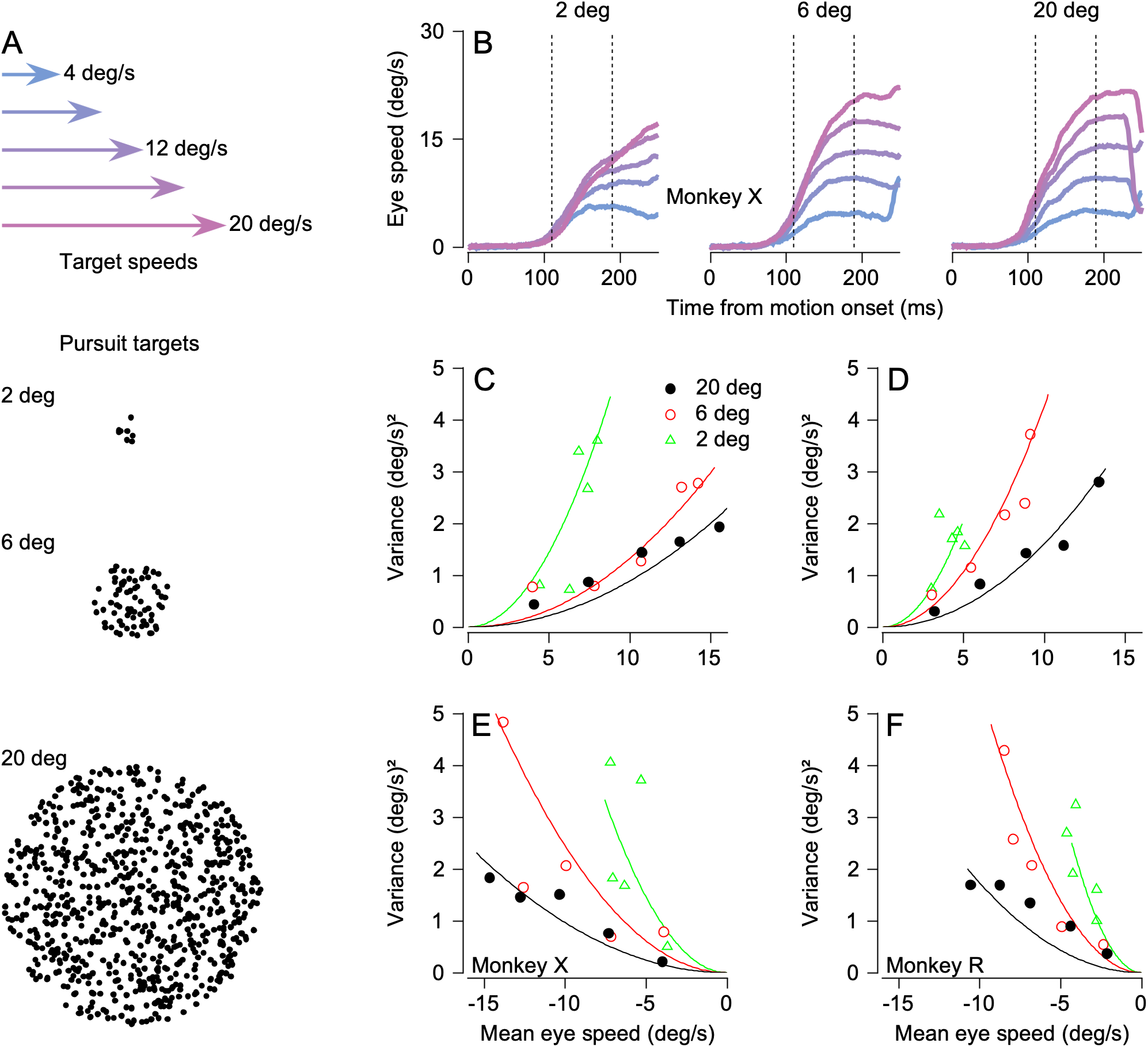
Target size affects the relationship between the variance and mean of pursuit initiation. A) Properties of pursuit stimuli. Top: Arrows indicate target speeds. Dot patches moved at a speed selected from a discrete uniform distribution with 5 values evenly spaced between 4 to 20 deg/s (colors). Bottom: Example dot patches corresponding to targets of three different sizes: 2 deg, 6 deg, and 20 deg. B) Mean pursuit initiation behavior for monkey X, sorted by target speed (colors; see arrows in panel A) and patch size indicated at the top of each graph. Each trace shows trial-averaged eye speed as a function of time, starting at the time of target motion onset. Vertical dashed lines indicate the window of time averaging used for subsequent analyses. C) Variance of eye speed of monkey X plotted against the mean eye speed for rightward pursuit. Symbols plot the behavior for different target speeds and sizes. Curves indicate the fit of a signal-dependent noise model where the Weber fraction is allowed to change with target size. D) As in panel C, but for rightward pursuit in monkey R. E) As in panel C, but for leftward pursuit in monkey X. F) As in panel C, but for leftward pursuit in monkey R. In C-F, green, red, and black symbols and curves show results for target sizes of 2, 6, and 20 deg, respectively.

Our new results show that the relationship between the variance and mean of pursuit responses depends strongly and unexpectedly on target size: changing target size disrupts signal-dependent noise. Figures 1C-F plot variance as a function of mean eye speed calculated from trial-by-trial, time-averaged data, using the interval from 110 to 190 ms after the onset of target motion (Figure 1B, vertical lines; Methods). For a given target size, variance increased with the mean eye speed for each individual target size. At a given mean eye speed generated in response to different target sizes, however, variance differed, breaking the relationship expected based on a standard interpretation of the psychophysical laws. Variance increases more rapidly as a function of mean pursuit initiation speed for the 2 deg patch (Figure 1C-F; green triangles) compared to the larger patches (red and black circles).

We use a signal-dependent noise model of the form *σ*^2^ = *w*^2^ *μ*^2^ to quantify the effect of target size on the relationship between mean, *μ*, and variance, *σ*^2^. The Weber fraction, *w*, captures the portionality between mean and variance and allows us to use changes in the Weber fraction to refer to the effect of target size on signal-dependent noise. We fitted the model to the data for 4, 12, and 20 deg/s patches and compared the predictions for the remaining two target speeds when the Weber fraction was fixed versus allowed to vary with target size. The model that allowed Weber fractions to vary (curves in Figure 1C-F) better predicted the variance associated with the 8 and 16 deg/s patches across patch sizes by allowing the Weber fraction to decrease with increasing target size (fixed *w* vs. flexible *w*: Monkey X, left pursuit: RMSE = 0.81 vs. 1.07 (deg/s)^2^; Monkey X, right pursuit: RMSE = 0.48 vs. 0.87 (deg/s)^2^; Monkey R, left pursuit: RMSE = 0.49 vs. 0.93 (deg/s)^2^; Monkey R, right pursuit: RMSE = 0.31 vs. 0.67 (deg/s)^2^). We observed the same decrease in Weber fractions with target size when we calculated *w* in 20 ms bins, with the additional (unsurprising, see^39^) finding that *w* decreases as a function of time across the initiation of pursuit (Supplementary Figure 1).

Intuitively, one might expect that the change in Weber fraction results from integrating the additional motion information associated with larger target sizes. However, this intuition is valid only when the biological sensors for motion improve their signal-to-noise as a function of target size or the downstream decoder can pool information from several sensors with independent sources of noise. Several experimental results argue against either scenario in the case of smooth pursuit. Motion sensitive neurons in area MT, the region of the primate cortex that contributes causally to pursuit^19–21^, exhibit decreased signal-to-noise as the size of a motion object extends beyond the classical receptive field^40, 41^, rather than the increased signal-to-noise required to explain our data. Further, individual MT neurons are subject to strong noise correlations^42, 43^, suggesting limits to the integration of motion information by downstream decoders of MT activity.

Neither are our results predicted by standard sensory or motor psychophysical models. Variance in the perception of speed is typically modeled as the result of signal-dependent noise in the sensory system under the assumption that sensory noise increases with speed, *s*, according to *σ*^2^ = *w*^2^*s*^2^, where *w* represents the Weber fraction for perception. However, Weber fractions for speed perception are constant across the range of target sizes used here^44, 45^, suggesting that the larger number of neurons activated by larger stimuli cannot be used to improve sensory signal-to-noise. Motor noise models represent variance as *σ*^2^ = *w*^2^ *μ*^2^, with *w* now representing a motor Weber fraction. As a result, behavioral variance is tied to the outgoing motor command^11^, and standard motor noise models do not predict any change in the Weber fraction for stimuli that lead to identical mean behavioral output.

### Target size affects the gain of visual-motor transmission

The fact that target size affects the average eye speed during the initiation of pursuit (Figure 1B) provides a clue to explanations for the effect of target size on pursuit Weber fractions. Figures 2B and C plot the eye speed of each trial, averaged across the interval from 110 to 190 ms following the onset of visual motion, as a function of target speed separately for each target size. The slope of the relationship between eye speed and target speed increased from 0.27 to 0.51 to 0.62 for the 2, 6, and 20 deg targets in monkey R, and from 0.45 to 0.78 to 0.81 in monkey X (95% confidence intervals were all less than ±0.025). For both monkeys, the slopes associated with the 6 deg target were significantly larger than those for the 2 deg target (monkey R: *z* = 36.16, *p* << 0.01; monkey X: *z* = 23.30, *p* << 0.01) and those associated with the 20 deg target were significantly larger than those for the 6 deg target (monkey R: *z* = 21.39, *p* << 0.01; monkey X: *z* = 3.37, *p* < 0.01).

**Figure 2.**
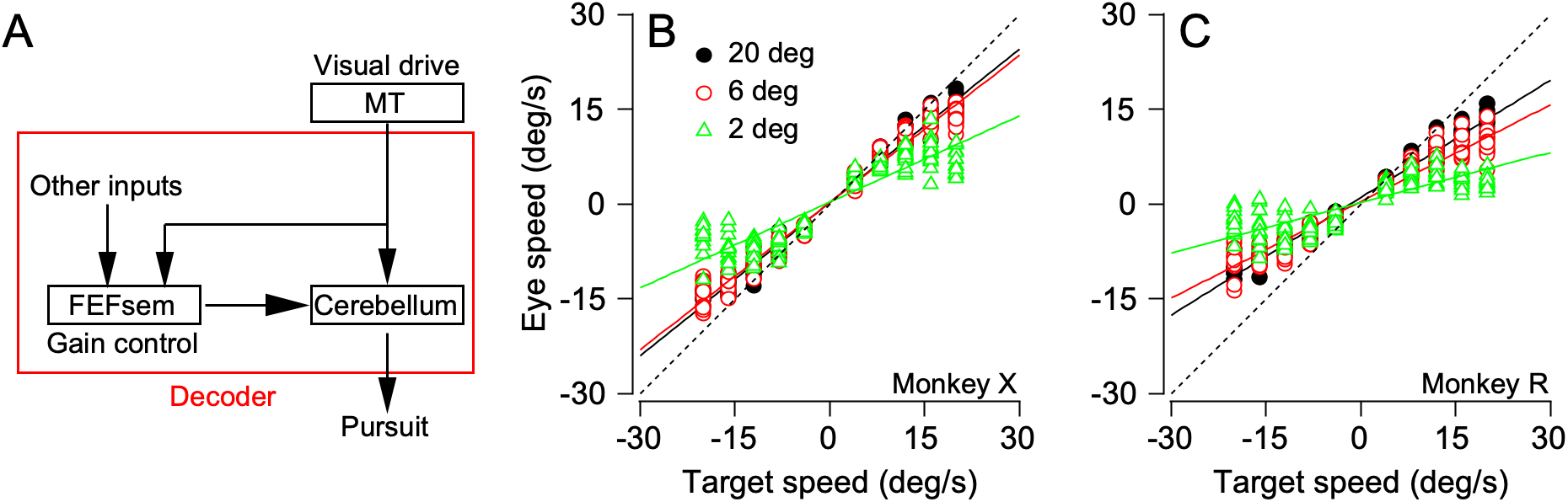
Gain of pursuit initiation increases with target size. A) Conceptual circuit model where neurons in area MT drive pursuit initiation through (at least) two parallel pathways: a direct pathway to the cerebellum and an indirect pathway for gain control through the smooth eye movement region of the frontal eye fields, FEF_sem_. Arrows indicate the direction of signal flow. The red box indicates that we can think of the entire pursuit circuit downstream of MT as the “sensory decoder”. B) Eye speed during pursuit initiation as a function of target speed for monkey X, sorted by patch size. Each symbol plots the average behavior for one combination of target speed and size. Lines plot the best fitting linear model to the 2 deg (green), 6 deg (red), and 20 deg (black) targets. C) As in panel B, but for monkey R.

Considerable prior research on pursuit eye movements allows us to think of eye speed (*E*) at the initiation of pursuit in simple terms as the product of two terms, *E* = *Gŝ*, where *ŝ* is an estimate of target speed based on sensory data, and *G* is the result of a process that controls the strength of visual-motor transmission^46^. Thus, the effect of target size on the slope of the relationships in Figures 2B and C could represent an effect of target size on *G* or *ŝ*. Available physiological evidence rules out the possibility that the change in slope arises from a change in the speed estimate, *ŝ*. The response of individual MT neurons to targets of different size has little to no effect on the speed tuning preferences of MT neurons^19, 47^. Therefore, estimates of *ŝ* derived by finding the preferred speed at the peak of the population response^23, 24, 48–50^should not shift as a function of target size, and the increasing slope cannot be explained by an effect of target size on *ŝ*.

We propose instead that the effect of target size on the slopes of the relationships in Figures 2B and C reflects an effect on the gain of visual-motor transmission, *G*. Therefore, we should focus on the neural circuits that control *G* to explain the companion effects of target size on the Weber fraction of pursuit. Our previous research on the initiation of pursuit has pointed to an overall neural circuit that has 2 parallel pathways (Figure 2A). A direct pathway estimates target speed (*ŝ*) from the population response in MT. An indirect pathway combines inputs from MT and other inputs to determine how strongly the motor system will respond to a given estimate of target speed by setting the value of *G*. Considerable evidence supports the existence of this second, “gain-control” pathway^27, 29^ and assigns this function to the smooth eye movement region of the frontal eye fields or FEF_sem_^26, 30^. According to everything we know, changing the size of the target should increase the total activity in the MT population response without shifting the peak of the population response, which should increase the the gain of visual-motor transmission (*G*) without changing *ŝ*, and cause the changes in slope in Figure 2B and C.

### Noise in the gain of visual-motor transmission captures the observed changes in Weber fractions

If changing target size alters pursuit initiation by controlling the strength of visual-motor transmission (*G*), then we should look at the pathways that control *G* as the potential basis for the companion changes in the Weber fraction. Indeed, the challenges of explaining the changes in the Weber fraction in terms of sensory or motor noise sources supports a source of noise in the sensory-motor decoder^51^.

To develop an intuition for how the pathway that controls visual-motor gain might also influence Weber fractions, we simplified the pursuit system into a computational model that is analytically tractable (Figure 3A). The model applies a gain, *G*, to the sensory estimate, *ŝ*, to generate motor output according to (*G* + *η_G_*)(*ŝ* + *η_s_*) + *η_m_*. The model includes three possible sources of behavioral variation:

1. Sensory noise, *η_s_*, which we model as Gaussian with variance that increases with sensory signal according to 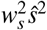 to accommodate the increase in perceptual variance associated with increased stimulus magnitude (e.g. Weber’s law^2, 44^).
2. Motor noise, *η_m_*, which increases with mean motor output, *μ*, according to 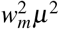^11, 52^.
3. Gain noise, *η_G_*, which is Gaussian with variance *σ*^2^.

**Figure 3.**
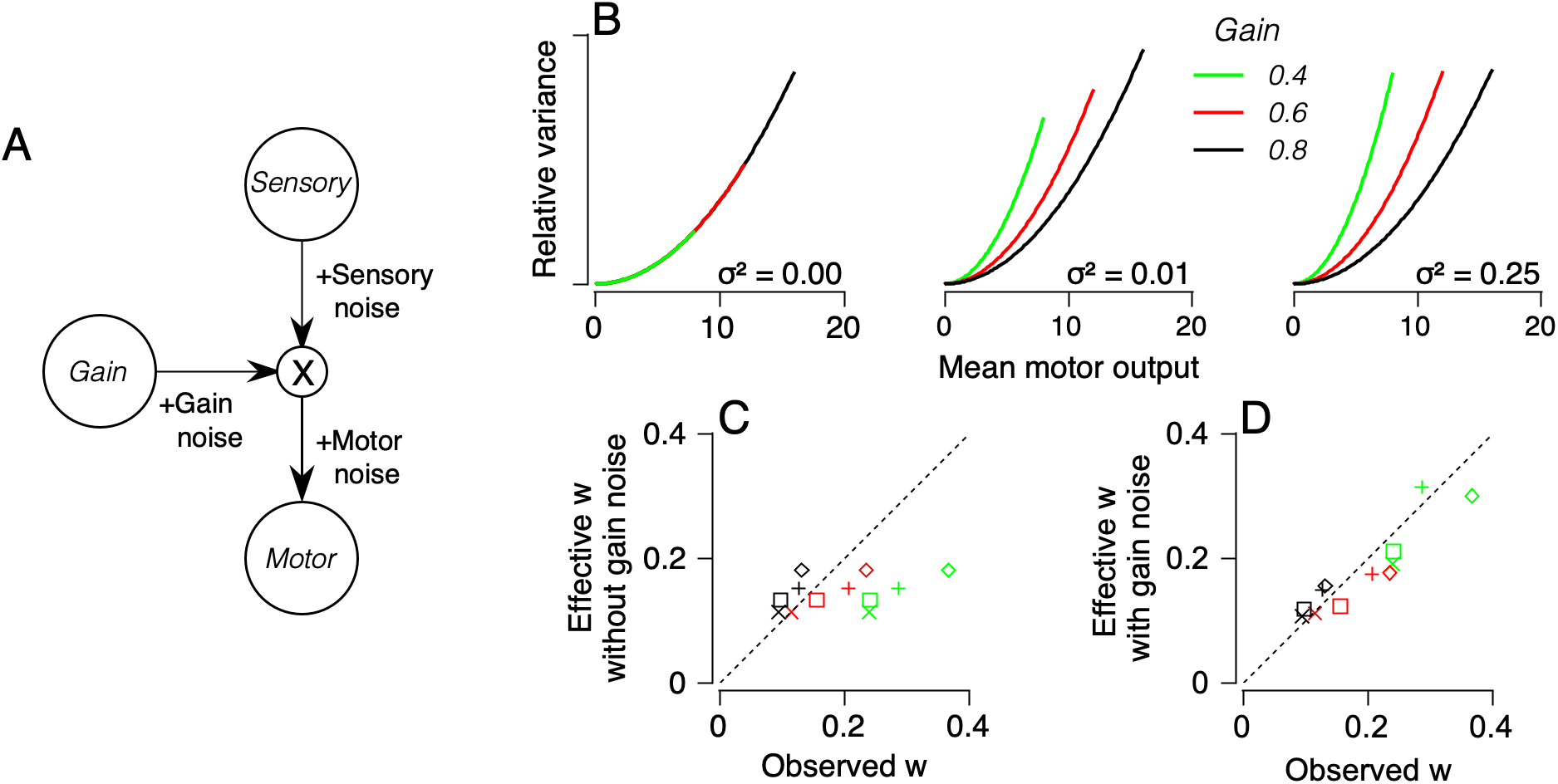
The “gain-noise” model: effect of gain noise on the relationship between variance and mean eye speed in behavior. Simplified model of sensory-motor transformations, where sensory input is transformed by the application of a gain signal to determine total motor output. B) Motor variance as a function of mean behavioral output for the simple model (Equation 1). From left to right, the three graphs show the predictions for gain noise with three different values of variance, *σ*^2^. Variance is plotted relative to the maximum across gains for each value of noise variance. Green, red, and black curves show predictions for three different values of gain at each value of noise variance. C) The effective value of the Weber fraction, *w*_eff_, inferred from a fit of the simple model without gain noise is plotted against the value of the Weber fraction fitted directly to the data for each target size separately (colors correspond to target size, as in Figure 1C). Plus signs and diamonds indicate right and left pursuit, respectively, for Monkey R. x’es and squares indicate right and left pursuit for Monkey X. Different colors represent different target sizes. D) As in panel C, but with gain noise included in the model.

If the sources of noise are statistically independent, then the mean output, *μ*, is *Gŝ* and its variance is 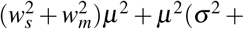 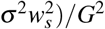. Therefore, this simple model produces the standard relationship between the mean and variance, *w*^2^ *μ*^2^, where the standard Weber fraction is replaced by the effective Weber fraction in the simple model:

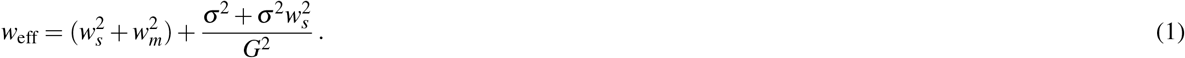

We can see from Equation (1) that the effective Weber fraction, *w*_eff_, will depend on *G* if (and only if) *σ*^2^ is non-zero.

Graphical analysis of Equation (1) confirms that the mean gain becomes an important factor in setting the value of *w* and therefore the relationship between variance and mean speed in the presence of gain noise (*σ*^2^ *>* 0). With gain noise, the variance of motor behavior increases with the mean motor output (*μ*) at a rate that depends on both the mean sensory-motor gain (*G*) and the variance of gain (*σ*^2^) (Figure 3B, middle and right graphs). The model predicts that the rate of increase in the variance of motor output will shift in a way that mimics our behavioral data. In the absence of gain noise (*σ*^2^ = 0), variance grows with mean motor output 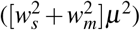 but the model predicts that Weber fractions will be constant across different levels of gain (Figure 3B, left). While the exact relationship between mean output and variance depends on the linearity of the model, the key prediction of a decreasing Weber fraction with increasing mean gain is robust to relaxing the linearity assumption (Supplementary Figure 2).

The simple model in Figure 3A fits the change in Weber fraction we observed in pursuit if and only if we allow gain noise. The effective Weber fraction for the model fitted with gain noise agreed with our behavioral observations across monkeys, directions, and target sizes (Figure 3D), whereas the effective Weber fraction for a model without gain noise did not (Figure 3C). We further compared the quality of fit of the model with versus without gain noise by fitting models to data from target motions at 4, 12, and 20 deg/s and testing on data from target motion at 8 and 16 deg/s. As before, a model with gain noise better predicted the data than a model that assumed no gain noise when fit to data (predicted variance with vs. without gain noise; Monkey X, left pursuit: RMSE = 1.18 vs. 1.54 (deg/s)^2^; Monkey X, right pursuit: RMSE = 0.81 vs. 0.96 (deg/s)^2^; Monkey R, left pursuit: RMSE = 0.96 vs. 1.29 (deg/s)^2^; Monkey R, right pursuit: RMSE = 0.68 vs. 0.94 (deg/s)^2^). Because there is no influence of the motor noise parameter, *w_m_*, on the changes in Weber fraction predicted by the model, we neglected *w_m_* in the analysis for Figure 3C and D (Methods).

### Biomimetic model for gain control in pursuit

The success of a model in predicting a single experimental result is not a strong test of the model, especially in an experimental system like smooth pursuit eye movements where so much is known about the statistics of neural and behavior responses. Therefore, we have elaborated the model by creating a biologically-realistic MT population response and processing it through a sensory decoder that includes the two known pathways that perform sensory estimation and control of visual-motor gain.

Our goal was for the elaborate, more “biomimetic” model to capture the following data:

1. The model’s signal dependent noise should shift with target size as in behavior.
2. The variance of the model’s output should match the overall variance of behavior.
3. The output of the model should exhibit trial-by-trial correlations with model MT neurons of the correct magnitude^24^.

We built a motion representation from model neurons that mimic the statistics of the spike counts of neurons recorded from area MT. Each model neuron was selective to a given direction and speed of motion, with mean tuning functions based on the well characterized properties of individual MT neurons and distributed in accord with published data^23, 43, 53, 54^ (Figure 4B). We distributed receptive field centers from 1 to 30 deg eccentricity based on the density of MT neurons per degree of visual angle^55, 56^, we set each neuron’s receptive field size based on the known relationship of size with eccentricity^57^, and we implemented inhibition outside the classical receptive field according to data from the literature^40, 41, 47, 58, 59^ (Methods; Figure 4D). We generated many single-trial population responses (Figure 4E) for each target speed and size by (1) measuring mean responses from the model population’s tuning curves, (2) adding noise from samples of a Poisson distribution, and (3) constraining the noise with neuron-neuron noise correlations as measured in MT^42, 43, 51^ (Figure 4C).

**Figure 4.**
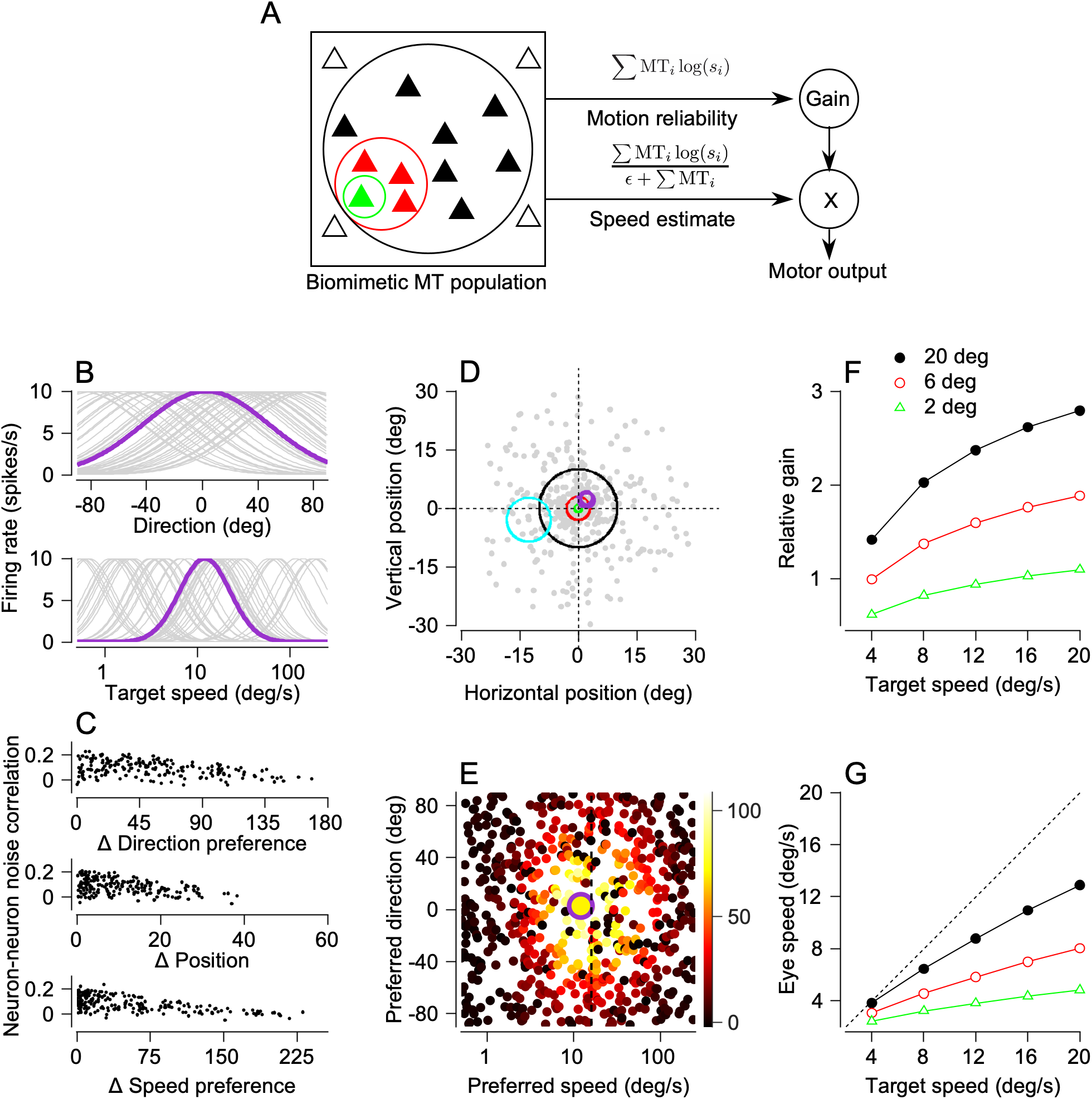
A more biomimetic model of MT and the downstream pursuit circuit. A) Overview of the biomimetic circuit model of pursuit. Activation of model MT neurons (triangles within square, left) depends on patch size (colored circles). Each neuron (MT_*i*_) drives (1) the motion reliability pathway, which computes a vector sum of MT activity (top) and (2) the speed estimation pathway, which computes a vector average (bottom). Application of the output of the motion reliability pathway to the output of the estimation pathway generates simulated pursuit (right). B) Direction tuning (top) and speed tuning (bottom) of a random sample of model MT neurons. Purple highlights one neuron. C) Noise correlations between model neurons as a function of the difference in direction preference (Δ Direction preference; top), distance between receptive field positions (Δ Position; middle), and difference in speed preference (Δ Speed preference; bottom). Each dot plots the measured noise correlation for a different pair of model neurons sampled from the population. D) Receptive field centers (gray points) of the population of MT neurons used to represent motion for pursuit. Green, red, and black circles correspond to the locations of the 2 deg, 6 deg, and 20 deg targets. Purple and cyan circles plot the classical receptive field of two example neurons. E) Response of each model neuron for an example trial (spikes/s; color). The center of each point corresponds to the neuron’s preferred direction and speed. Large purple symbol indicates the example neuron from panels B and D. F) Mean gain of the population as a function of target speed and size (colors as in panel A). G) Mean simulated eye speed as a function of target speed and size (colors as in panel A).

We constructed a sensory decoder that transmits the output of each model MT neuron along two pathways (Figure 4A), an architecture that accounts for a considerable amount of published behavioral and physiological data^30, 60^. The upper pathway performs vector summation of model MT neural responses, following previous results relating motion reliability to the amplitude of the MT population response. It uses that amplitude to set the gain of sensory-motor transmission, *G*^30^. The lower pathway performs vector averaging of the MT population response to estimate target speed, *ŝ*^24, 48, 50^. We then multiply the outputs of the two pathways (*Gŝ*) to complete decoding and generate simulated pursuit behavior. Both of the pathways are needed, for example, to account for the joint effects of target speed and contrast on eye speed in the initiation of pursuit^30, 61^.

Because larger motion patches increased the number of model MT neurons activated by the stimulus (Figure 4D), vector summation performed by the motion reliability pathway generated a gain that increased with patch size when averaged across trials (Figure 4F). Gain also increased with target speed due to the speed tuning properties of MT neurons (Figure 4F). Multiplication of the output of the gain pathway times the target speed provided by the vector averaging speed estimation pathway resulted in mean eye speeds similar to those observed in our monkeys (Figure 4G). Specifically, the decoder’s output generated mean eye speeds that increased steeply as a function of target speed at a rate that depended on target size.

### Noise is required in the gain-control pathway to capture the statistics of behavioral and neural data

When we add noise in the gain-control pathway, our biomimetic model predicts all 3 of the key statistics we set out to reproduce: an effect of target size on Weber fractions, realistic magnitudes of variance in eye speed at the initiation of pursuit, and realistic trial-by-trial correlations between the activity of individual MT neurons and pursuit eye speed. Without noise in the gain-control pathway, the model failed to predict any of these statistics.

Comparison of Figures 5B and C shows that noise in the gain-control pathway allows the rate of increase in the modeled variance to depend on the size of the target: the variance in the simulated response to the 2 deg patch (green triangles) increases more rapidly than for the 6 deg patch (red circles), with a still slower rise for the 20 deg patch (black circles). The presence of gain noise also increases the magnitude of the variance in eye speed so that it matches more closely the values in the data from our monkeys: note that the y-axes are the same in Figures 1C-F and 5B-C. Across a wide range of parameterizations of our circuit model we found no instantiation of the circuit that could both recapitulate the effect of target size on Weber fractions and predict realistic amplitudes of behavioral variance without the inclusion of noise in the motion reliability pathway (Supplementary Figure 3). Increased sensory noise and/or increased magnitudes of noise correlations in our model MT neurons would increase the magnitude of eye speed variance, but only by contradicting available physiological data from MT^43^.

**Figure 5.**
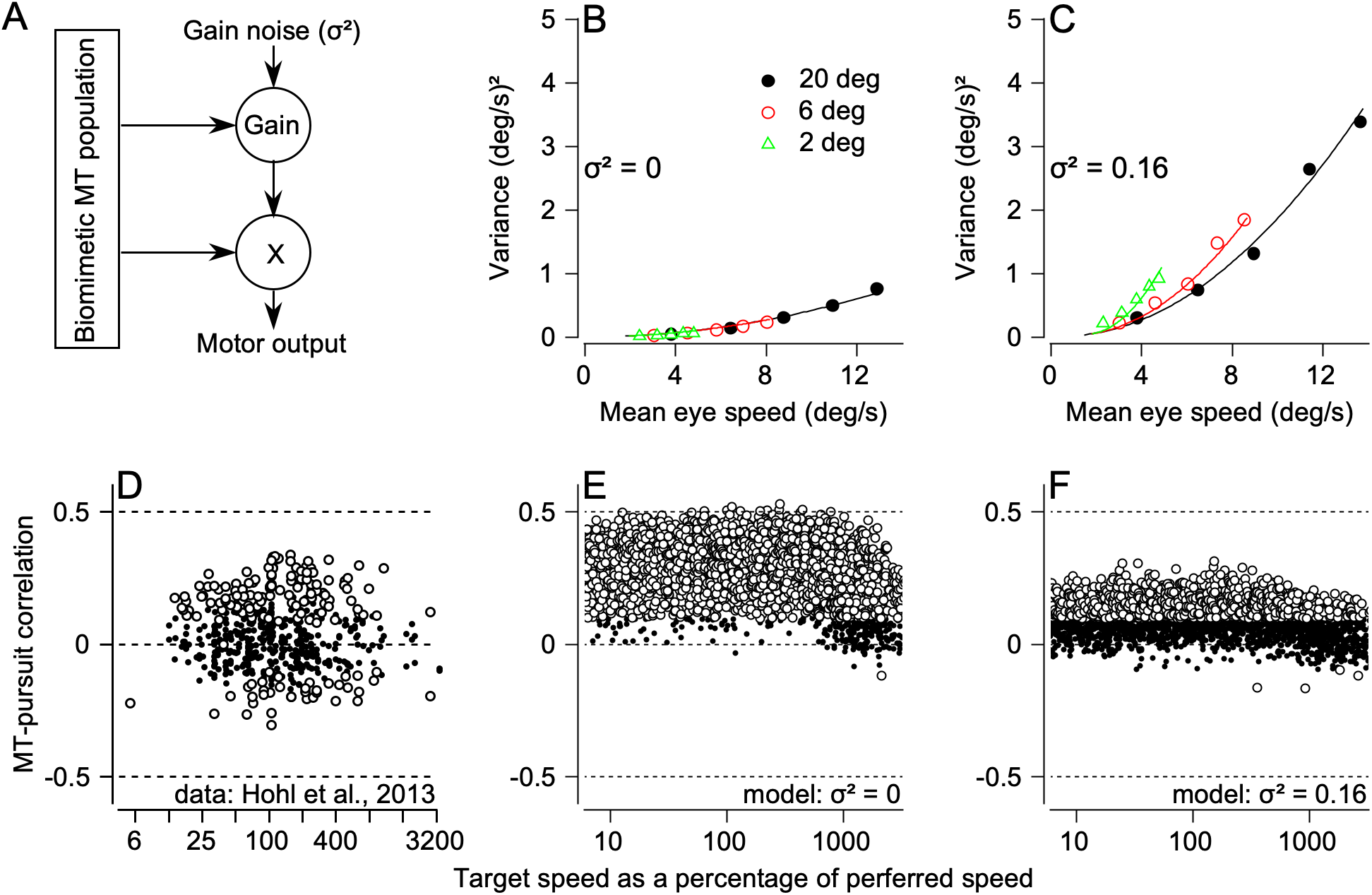
Biomimetic model with gain noise captures key features of behavioral and neurophysiological data. A) Biomimetic model as in Figure 4A, but highlighting noise added to the gain pathway. Noise was drawn from a zero-mean Gaussian with variance *σ*^2^. B) Relationship between variance and mean eye speed of model output without gain noise. Green triangles, red circles, and black circles correspond to model behavior in response to 2, 6, and 20 deg targets, respectively. Green, red, and black curves plot fits of the gain-noise model from Equation (1) to circuit model responses to 2, 6, and 20 deg targets, respectively. C) As in panel B, but with gain noise. D) Observed MT-pursuit correlations, reproduced with permission from^24^. Each symbol plots the trial-by-trial neuron-behavior correlation for an individual neuron. Open circles indicate statistically significant correlations (p<0.05). E) MT-pursuit correlations measured for the model neurons in the biomimetic model without gain noise when the patch size was 6 deg. Conventions as in panel D. F) As in panel E, but with gain noise.

Comparison of Figures 5D-F shows that noise in the gain pathway allows the model to reproduce the MT-pursuit correlations recorded by Hohl et al. (2013)^24^. To obtain the data in Figure 5D, the authors recorded up to 500 responses of individual MT neurons to the same moving target during the initiation of pursuit. They then computed trial-by-trial correlations between the spike count of the MT neuron and the eye speed in the initiation of pursuit and plotted the result as a function of the relationship between the target speed and the preferred speed of the neuron. They found statistically significant, mostly positive correlations in many neurons (Figure 5D).

Without gain noise (Figure 5E), the model predicts MT-pursuit correlations that are mainly positive, as in the data, but the magnitude of the correlations is considerably larger than in the data (Figure 5D). Gain noise is effectively “noise added downstream” and therefore reduces the magnitude of the model’s MT-pursuit correlations (Figure 5F) to a level closer to the measured values. We also note that the 2-pathway decoder used in our model (Figure 5A), reproduces the unexpected finding the MT-pursuit correlations were positive whether the target speed was faster or slower than the preferred speed of the MT neuron, a fact that Hohl et al. (2013) struggled to explain^24^. Overall, the presence of gain noise has the expected effect of adding downstream noise. It both increases the variance of model output and partially decorrelates the response of MT neurons from model output^15^.

## Discussion

Behavior results from decoding sensory representations. There are many examples where application of decoding equations to sensory representations have elucidated how the brain generates motor or perceptual behavior^3–8^. However, to accommodate the complexity of the natural behavioral repertoire, a decoder requires the capacity to flexibly map sensory inputs to motor outputs. By incorporating the known structure of the neural system for smooth pursuit into a decoder and leveraging structural noise, we have created a decoder that is biologically more realistic and that reproduces the complete statistics of sensory-motor behavior more powerfully than traditional decoding equations. Our analysis demonstrates how the conventional decoding approach to understanding the neural basis for perceptual or motor behavior can be re-imagined and aligned better with biological reality.

We have combined theory, measurements of the statistics of behavior and neural responses, and explicit modeling of the functional pathways of the sensory-motor decoder. Our approach was driven by extensive knowledge of the relationship between the functional components of visually-guided smooth pursuit eye movements, and firing properties of neurons in the relevant circuits. Recordings made in monkeys during pursuit eye movements have led to a strong hypothesis for the structure of this sensory-motor system^30, 46^ and focused us on a sensory decoder with two pathways and two functional components. Both pathways rely on inputs from the sensory representation of visual motion in area MT. The first pathway estimates target speed, regardless of target size, by performing vector averaging of the MT population response. Vector average decoders successfully capture much of pursuit behavior^4, 24, 43, 50^, and have the critical emergent property that they predict larger magnitude pursuit initiation for faster target speeds. The second pathway computes the gain of the sensory-motor transmission based on the fact that the evoked eye movement is modulated by the context of a visual motion input^27–29^. The responses of neurons in FEF_sem_ substantiate the 2-pathway architecture and the inclusion of gain control^62–64^ and FEF_sem_ has a causal relationship with the gain of visual-motor transmission^26, 65, 66^. Although made quite specific here, our results apply generally to decoders of any architecture that implements the computation *f*(**r**_MT_)**w**^*T*^ **r**_MT_, where **w** is a vector of decoding weights, **r**_MT_ is a vector of sensory responses, and *f* (**r**_MT_) returns a gain with noise that is independent of **r**_MT_.

Building a two pathway structure into our sensory decoder was an essential prerequisite for capturing our observation that changing target size also changed the Weber fraction for the initiation of pursuit. The two-pathway structure allows for a plausible source of independent noise in gain control that captures the changes in Weber fraction we observe. The resulting model produced output that reproduced the measured amplitude of behavioral variation, and also matched the trial-by-trial correlations between model MT responses and pursuit behavior to those observed in physiology. Thus, a decoder with multiple functional pathways and a finite contribution of noise provides a parsimonious account of a wide-ranging set of results.

Previous publications have concluded that the variation in sensory-motor behavior could be explained in terms of noise in the sensory system alone for both pursuit and other motor behaviors^67, 68^. However, increasing evidence from both behavior^16–18^ and physiology^14, 69, 70^ suggests that neural systems between sensation and action, including gain systems^31, 71^, are subject to noise that propagates into motor output. Under standard experimental conditions, the expected impact of gain noise on behavioral variation is identical to that of sensory noise and so the analytical results of Osborne et. al. (2005)^67^ would emerge whether behavioral variation arises from sensory noise only, or from a combination of sensory and gain noise. Thus, our conclusion, that the computation of sensory-motor gain is subject to an independent source of noise that arises from gain control, does not contradict previous data, only previous conclusions.

Several lines of evidence argue against the logical idea that integration of the additional information associated with increasing target size should decrease the effective Weber fraction in behavior^72^, and therefore explain our behavioral results. First, due to surround suppression^40^, the average MT neuron exhibits decreased precision as target size increases beyond the classical receptive field^41^, not the increased precision that would be required to explain our data. Second, increasing target size does not lead to increased precision during perceptual decision making that is based on the same sensory representation that drives pursuit initiation^44, 45^. Finally, the correlated variability of MT neurons appears to limit the increase in information with increasing population size^43, 51^. Indeed, our model’s inclusion of correlated variability in MT neurons with different receptive field locations provides a parsimonious account of results from experiments with both motor and perceptual endpoints.

MT neurons drive the initiation of pursuit^19–21^. Therefore, it was important in creating our model MT population response to assess critically how stimuli outside the classical receptive fields of MT neurons influence the representation of motion by the population. We modeled our neurons with a uniform surround that divisively inhibits the response of the classical receptive field^40, 41, 50^. Under this assumption, made because surround stimulation changes the amplitude but not other properties of MT neuron speed tuning^47^, our decoding model is robust to the degree of suppression by surround stimulation. However, extra-classical surrounds of MT neurons are varied and complex^73^, and a still more accurate model of how our stimuli impact the responses of MT neurons and downstream systems would be the next step to completely explain the complex spatial integration properties of pursuit^74^.

Decoding architectures that employ gain control apply broadly to sensory-motor behaviors. Gain adjustment is a critical feature of optimal estimation by Bayesian integration, where the weight given to a sensory cue depends on the relative reliability of each potential source of information. Reliability-based weighting strategies operate when subjects integrate sensory cues^72, 75–79^ or combine measurements with prior knowledge based on past experience^29, 33, 80, 81^. Therefore, neural mechanisms must exist for appropriately applying sensory gain during estimation. Gain control also plays a role in control strategies to optimize motor behavior^82, 83^. Here, the flexible application of gain on sensory estimates modulates control efforts that are relevant to completing the task^12, 34^. Given the importance of gain control to sensory-motor behaviors, it is perhaps not surprising that dedicated pathways exist in neural circuits to learn and implement context-specific sensory-motor gains^30^. Our findings suggest that the flexibility afforded by independent gain control pathways comes at a cost of increased variability associated with noise that arises through imperfect circuit computations.

More generally, our results have broad applicability to models that attempt to explain signal and noise in complex behaviors. The brain’s computational power is achieved through numerous pathways that process information in parallel before ultimately driving motor output. Our results stress the importance of understanding how noise in each of the processing pathways contributes to behavioral variation. If we can characterize and model the pathways in sensory decoders for movement and identify noise specific to each neural pathway, then we can constrain the possible circuit implementations and generate specific, experimentally-testable predictions for circuit function.

## Methods

Two adult, male rhesus macaques weighting between 12.0 and 16.8 kg performed the pursuit task. All experimental protocols were approved by the *Institutional Animal Care and Use Committee* at Duke University and performed in accordance with the National Institutes of Health *Guide for the Care and Use of Laboratory Animals*. Eye movements were tracked with a scleral search coil while their heads were fixed by a restraint post^84^. Implantation of experimental apparati was performed using sterile procedures under general anaesthesia with isoflurane. After implantation, animals were trained to fixate and pursue visual targets for juice reward based on the analog signal from the implanted search coil. Both animals had extensive experience performing pursuit tasks before this experiment.

### Stimuli and task

Visual targets were displayed on a 23” CRT monitor with an 80 Hz refresh rate at a distance of 30 cm. At this distance the screen subtended 77 deg of visual angle horizontally and 55 deg vertically. Stimulus presentation and timing were controlled by custom software (Maestro).

Each trial began with the monkey fixating a white, 0.5 deg, circular fixation point for a random interval selected from a uniform distribution between 300 to 400 ms. After the monkey maintained fixation within 3.5 deg for the entire interval, the fixation point disappeared and was replaced by a target for pursuit. Targets were dot patches presented in circular aperture with a diameter selected each trial uniformly from one of three values: 2, 6, or 20 deg. All targets were 100% contrast, white dots on a black screen, with dot density set to 2.55 dots/deg^2^ and the placement of the dots randomized across trials. Upon presentation, dot patches moved either to the right or left within their aperture for 150 ms before continuing to move globally across the screen for 750. This procedure reduced the occurrence of catch up saccades during pursuit initiation^22^. Patches moved at a speed selected for each trial from a discrete, uniform distribution with 5 speeds linearly spaced from 4 to 20 deg/s. Monkeys were rewarded for maintaining their eyes within 3.8, 3.8, or 10 deg of the center of the visual target throughout the presentation of the 2, 6, and 20 deg diameter targets, respectively; trials were aborted if the monkey’s eye strayed outside of this window. After either successful completion or an aborted trial, the fixation point for the next trial was presented immediately.

### Data analysis

Horizontal and vertical eye position and speed for each trial was stored for offline analysis. Monkey R and monkey X completed 4027 and 1442 total trials, respectively. For each trial, we first detected the presence of saccades within a 250 ms window starting from motion onset. Any trial with a saccade was discarded from future analysis. After discarding these trials, we averaged the horizontal speed on each trial within a window 110 to 190 ms relative to target motion onset and then computed statistics across all trials with identical targets and target motions. Subsequent analyses were performed using the time-averaged data.

To test if Weber fractions associated with the initiation of pursuit changed with target size we determined the mean and variance of pursuit speeds, conditioned on target speed, patch size, and motion direction. We then fit the behavioral data to two models to each motion direction, both of the form 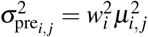, where *μ_i,j_* was the mean of the observed behavior, *i* indexes target sizes, and *j* indexes target speeds. Under the null hypothesis we held *w_i_* fixed across target size. Under the hypothesis that Weber fractions changed, we allowed *w_i_* to vary across target sizes. For each model, we minimized the following equation:

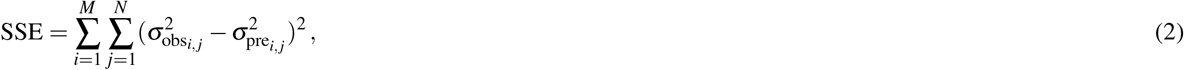

where *M* is the number of different patch sizes in a training set, *N* is the number of different target speeds in a training set, 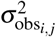 is the observed variance for the *i*th patch size and *j*th target speed, and 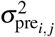 is the corresponding variance predicted by either hypothesis. For each model we minimized the sum of squared errors across all patch sizes and target speeds of 4, 12, and 20 deg/s. To test which model better fit the data, we computed the root mean squared error (RMSE) in the predicted variance of each model across patch sizes for target speeds of 8 and 16 deg/s. RMSE was defined as

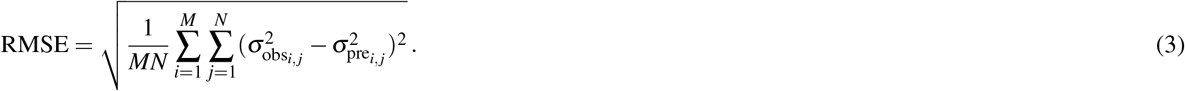

We performed a similar analysis to test if the data was consistent with gain noise. We fit two models to the data: the gain noise hypothesis, H_*σ>*0_, or the null hypothesis without gain noise, H_*σ*=0_. For each model, we minimized Equation (2), except we set

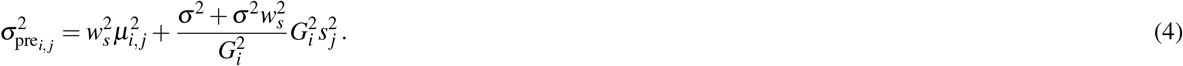

Because the term *G_i_* cancels, *w_s_* and *σ* are the only free parameters for the gain noise hypothesis. For the null hypothesis, *w_s_* was a free parameter and *σ* was set to 0. We followed the same procedure as above, fitting the data across patch sizes and target speeds of 4, 12, and 20 deg/s and testing the fit by calculating the RMSE according to Equation (3) across patch sizes for target speeds of 8 and 16 deg/s. To estimate effective Weber fractions under the gain noise model, we used the fit values of *w_s_* and *σ* for each model, and *G_i_* was found via linear regression of the eye speed vs. target speed data (i.e. Figure 2B and C).

### Biomimetic circuit model of pursuit

Our circuit model of pursuit was based on a population of model MT neurons with realistic tuning properties. Based on estimates of the cortical size of the representation as a function of eccentricity in mm^2^ and the density of cortical neurons per mm^255, 56^, we randomly sampled receptive field centers spanning eccentricities of 1 deg to 30 deg. Based on this empirical evidence we estimated that there would be a total of 667 MT neurons within this range, but our results do not critically depend on the exact number of MT neurons simulated (Supplementary Figure 4).

We modeled the response to a target of direction *θ* of each MT neuron, indexed by *i*, as

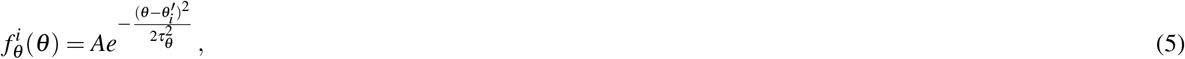

where *A* is the maximum amplitude of the direction response, 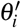 is the preferred direction, and *τ_θ_* determines the width of the response function. We set *A* = 10 and *τ_θ_* = 45. The preferred direction was sampled from a uniform distribution between −90 and 90.

We modeled the response of each model neuron in response to a target of speed *s* as

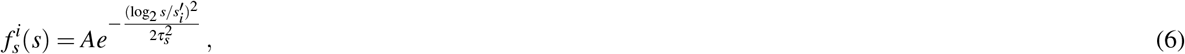

where *A* is the maximum amplitude of the speed response, 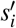 is the preferred speed, and *τ_s_* determines the width of the speed response in logarithmic space. We set *A* = 10 and *τ_s_* = 1.64. The preferred speed was sampled from a uniform distribution between 0.5 and 256 deg/s.

Finally, we modulated the response based on the overlap the dot stimulus had with the receptive field of each neuron. Based on previous measurements^57^, we assumed the receptive field diameter grew with the eccentricity according to 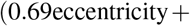 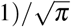. We then determined the response based on stimulation of the classical receptive field as

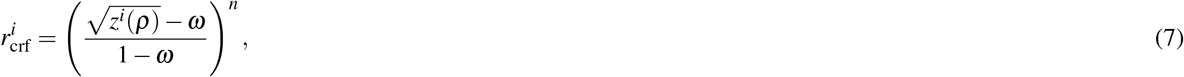

where *z^i^* takes a value between 0 and 1 that represents the fraction of the classical receptive field overlapped by the stimulus of size *ρ*. *ω* and *n* model threshold nonlinearities in receptive field summation by MT neurons^85, 86^. Values of *n* near 1 will produce responses that are close to receptive field summation. Values approaching 0 allow the response of a neuron to approach its maximum whenever the stimulus partially overlaps with the classical receptive field.

Previous experiments have further documented that stimuli outside the classical receptive field of MT neurons tend to suppress the response to stimuli within the receptive field^40, 73, 87^. We assumed an extra classical surround with a radius three times the size of the classical receptive field. After calculating the overlap of the stimulus with the extra classical surround, 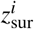, the response of the surround was

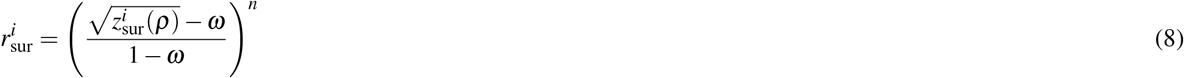

with the threshold nonlinearity identical to that used for the classical receptive field. We then modeled surround suppression as divisive normalization such that the response due to the receptive field properties was

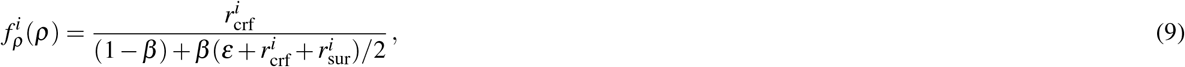

where *β* determines the degree of surround suppression, and *ε* is a constant which normalizes the response to be between 0 and 1 when *β* = 1. For each of the figures presented in the main manuscript, we set *ω* = 0, *n* = 1, and *β* = 1.

Putting together the response due to direction, speed, and receptive field position, the deterministic response of each neuron was then modeled as

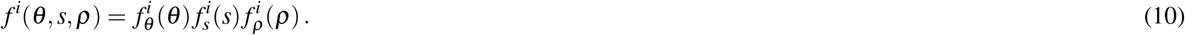

To emulate the noise properties of MT neurons, we added Poisson noise that was correlated across neurons. Following^43^, we modeled the correlated noise as

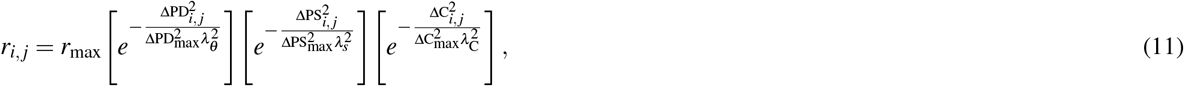

where i and j index neurons, *r_i,j_* is the noise correlation between the pair, *r*_max_ set the maximum value of the correlation, ΔPD_*i,j*_ is the difference in preferred motion direction, ΔPS_*i,j*_ is the difference in preferred speed, ΔC_*i,j*_ is the distance between receptive field centers, ΔPD_max_ is the maximum difference in preferred directions, ΔPS_max_ is the maximum difference in preferred speeds, ΔC_max_ is the maximum distance between receptive fields. *λ_θ_*, *λ_s_*, and *λ*_C_ set the rate of decay in correlation with increasing difference in preferred distance, speed, and receptive field position, respectively. We set *λ_θ_*, *λ_s_*, and *λ*_C_ to 0.40, 0.30, and 0.30, respectively and *r*_max_ to 0.36; all values based on those measured from physiology^43^. Correlated Poisson noise, 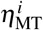, was then introduced as in^51^ to generate variable responses for each trial according to

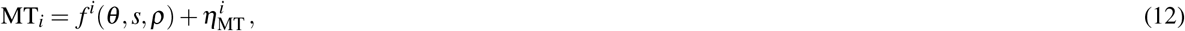

where MT_*i*_ is the firing rate of the *i*th model MT neuron.

Simulated pursuit initiation responses were generated according to the following steps. Activity from the population of model MT neurons was read off according to two pathways. The first calculated an estimate of the pursuit speed according to

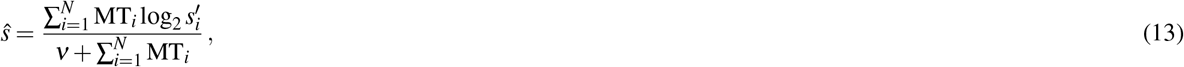

where *ν* was set to 0.05. The second set the gain according to

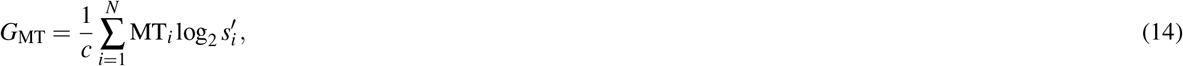

where *c* was set to make the mean pursuit, across speeds, for the 20 deg patch size equal to the mean across *s* of log_2_ *s*. The simulated pursuit speed, *m*, for each trial was then set to

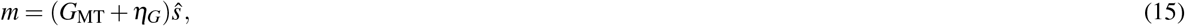

where *η_G_* was selected each trial from a Gaussian distribution with mean 0 and variance *σ*^2^. Subsequent analysis of the simulated pursuit responses followed that for the actual pursuit responses.

## Data availability

Data is available from the authors upon reasonable request.

## Code availability

Code is available from the authors upon reasonable request.

## Acknowledgements

This work was supported by NIH grant EY027373. We thank Stuart Behling, Timothy R. Darlington, David J. Herzfeld, Nathan J. Hall, Bing Liu, and Leslie C. Osborne for their comments on an earlier version of this manuscript. We also thank Stefanie Tokiyama and Bonnie Bowell for animal assistance.

## Author contributions

SWE and SGL devised the experiment. SWE derived predictions from the simple model. SWE and SGL conceived of the biomimetic model. SWE performed experiments and analyses. SWE and SGL wrote the manuscript.

## Additional information

### Competing interests

The authors declare no competing interests.

## Supplementary Materials

**Supplementary Figure 1.**
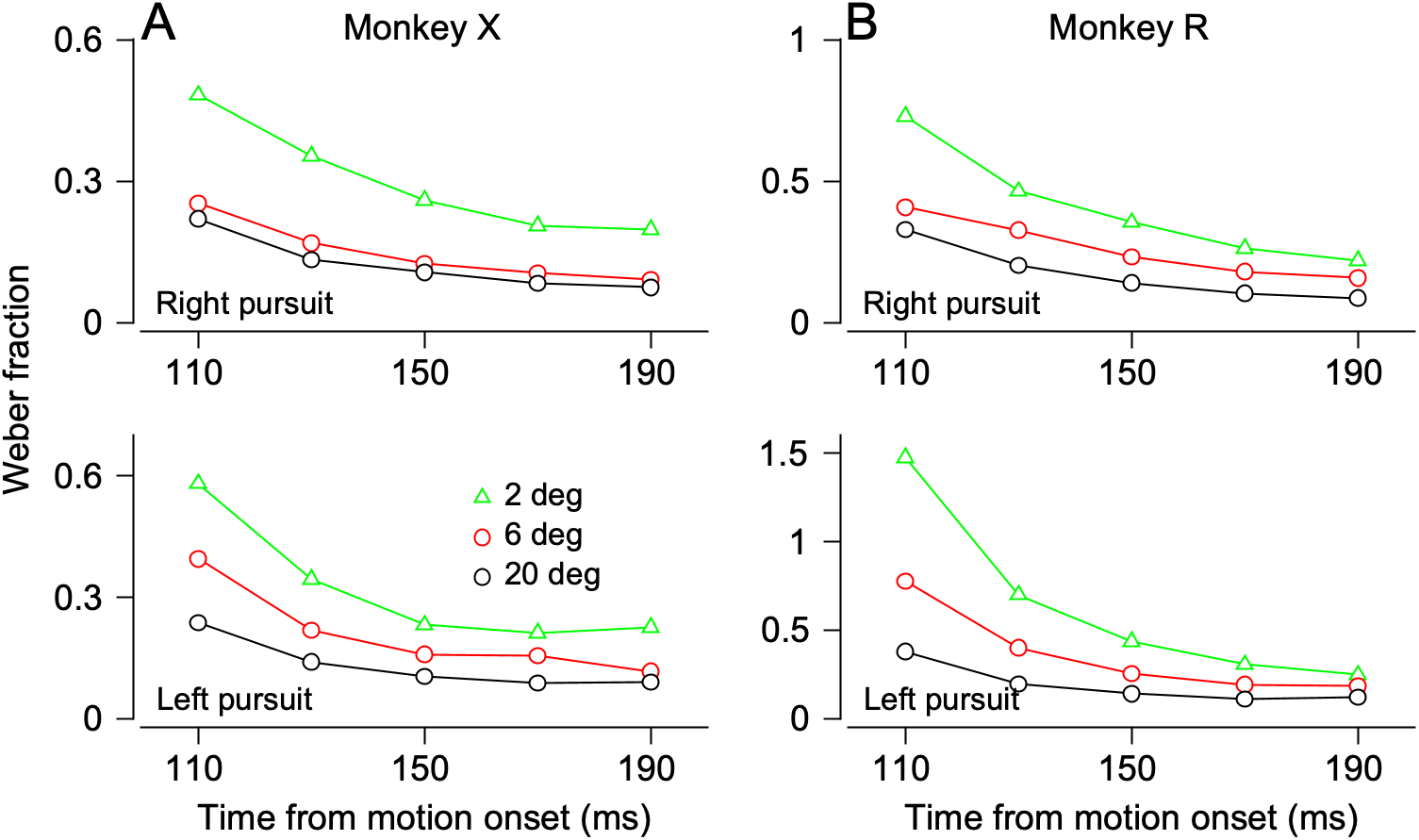
Target size affects Weber fractions across the full interval of the initiation of pursuit. We divided the interval from 110 to 190 ms after the onset of target motion into intervals of duration 20 ms and analyzed the Weber fraction for each target size in each interval. Green, red, and black symbols show data for target sizes of 2, 6, and 20 deg and indicate that the Weber fraction is consistently smaller for larger targets across the entire interval of pursuit initiation. Weber fraction declines for each curve across the initiation of pursuit, indicating that the mean eye speed increases more rapidly across time than does the variance of eye speed.

**Supplementary Figure 2.**
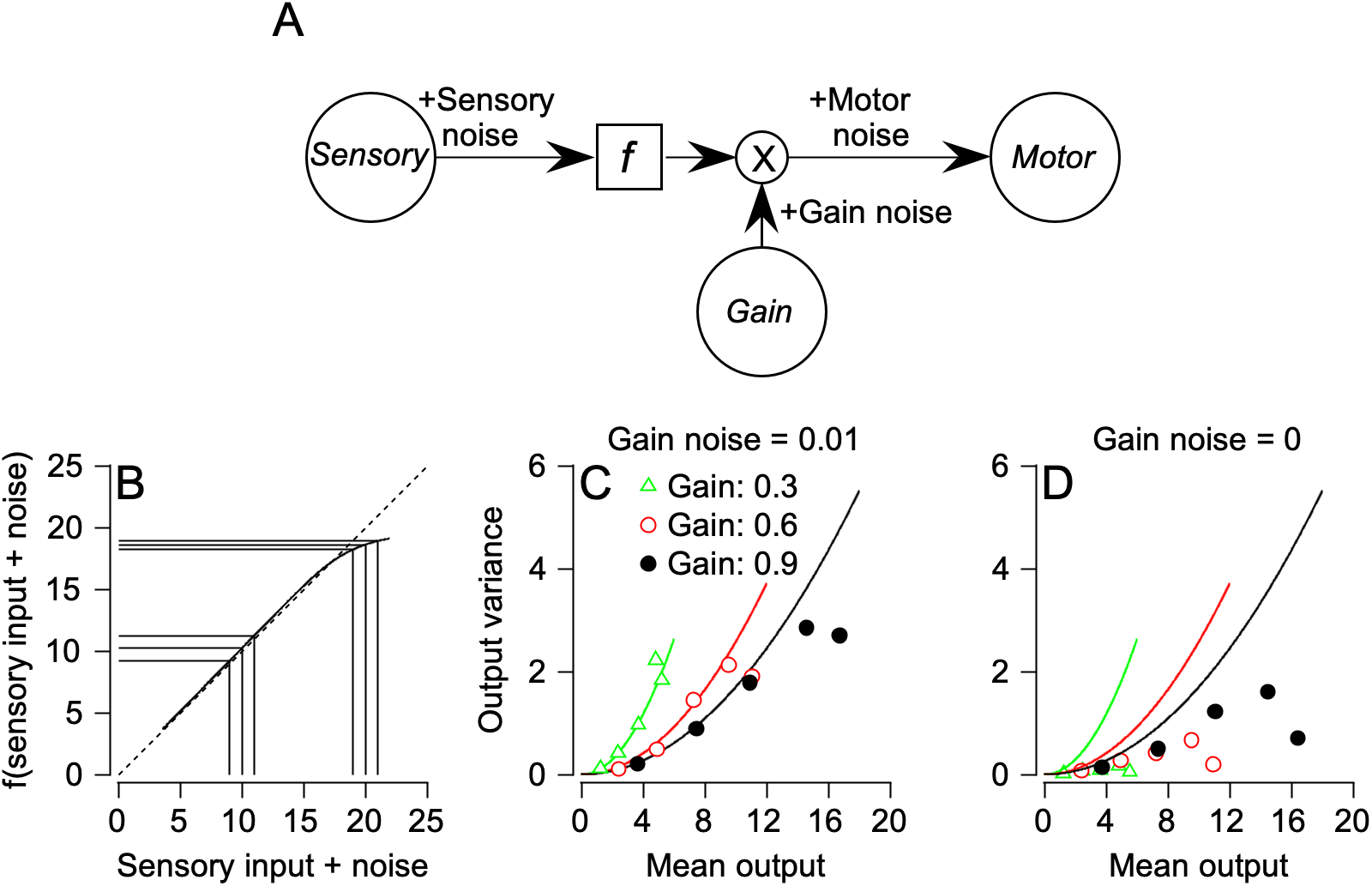
Generalization of the gain-noise model to nonlinear sensory encoding and estimation. To examine whether the signature of gain noise could result from other changes in model configuration, we generalized the simple model in Figure 3 to include a nonlinear transformation. In panel A, a sensory input, *s*, with signal-dependent noise, *η_s_*, is rendered non-linear according to the function *f*. A gain signal with mean *G* and variance *σ*^2^ is applied following the transformation by *f*, and motor noise added following the gain. For small values of *η_s_*, the variance will be:

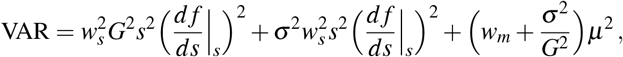

where *w_s_* is the Weber fraction that determines the magnitude of sensory noise according to target speed 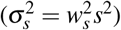, *w_m_* is the motor Weber fraction that determines the magnitude of motor noise according to 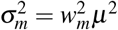, and *μ* is the mean output of the model. The first two terms of the equation cause the variance of motor output to depend on the derivative of *f*, *df/ds*, in the vicinity of *s*. The last term of the equation causes the output variance to depend on the mean gain when gain noise is non-zero. Panel B illustrates the effect of a nonlinear transformation, *f*, based on the Bayesian least squares (BLS) estimator for signal-dependent sensory noise (*w* = 0.1)^9, 72, 88^. In the vicinity of *s* + *η_s_* = 10, df/ds is near 1. As a result, sensory noise illustrated by the 3 parallel vertical lines near *s* + *η_s_* = 10, adds variance to the output of *f*, as shown by the three horizontal lines that project the sensory input onto the y-axis. In the vicinity of *s* + *η_s_* = 20, *df/ds* starts to approach 0. As a result the same amount of sensory noise shown by the vertical lines near *s* + *η_s_* = 20 leads to very little output variance, shown by the projection onto the y-axis (horizontal lines). Panels C and D plot the output variance as a function of the mean output amplitude for different configurations of the model and illustrate that a plausible non-linearity alone is not enough to produce the predictions of gain noise. Panel C reproduces Figure 1 with gain set to 0.3 (green), 0.6 (red), or 0.9 (black) and the gain noise set to have a variance of 0.01 (standard deviation of 0.1). The continuous curves show the best fits to the model outputs based on the model in Figure 3. Panel D illustrates the performance of the non-linear model with gain noise set to zero and shows that the nonlinearity does not convey the same effect as gain noise. Output variance is still signal-dependent, but the data do not separate according to the value of gain. Note that the nonlinearity should not affect the results for target speed in the linear range of *f*, but it might (but does not) cause the signature of gain noise for the highest values of target speed. The continuous curves in panel D are the same as those in panel C, for easy visual comparison of the performance of different configurations of the model.

**Supplementary Figure 3.**
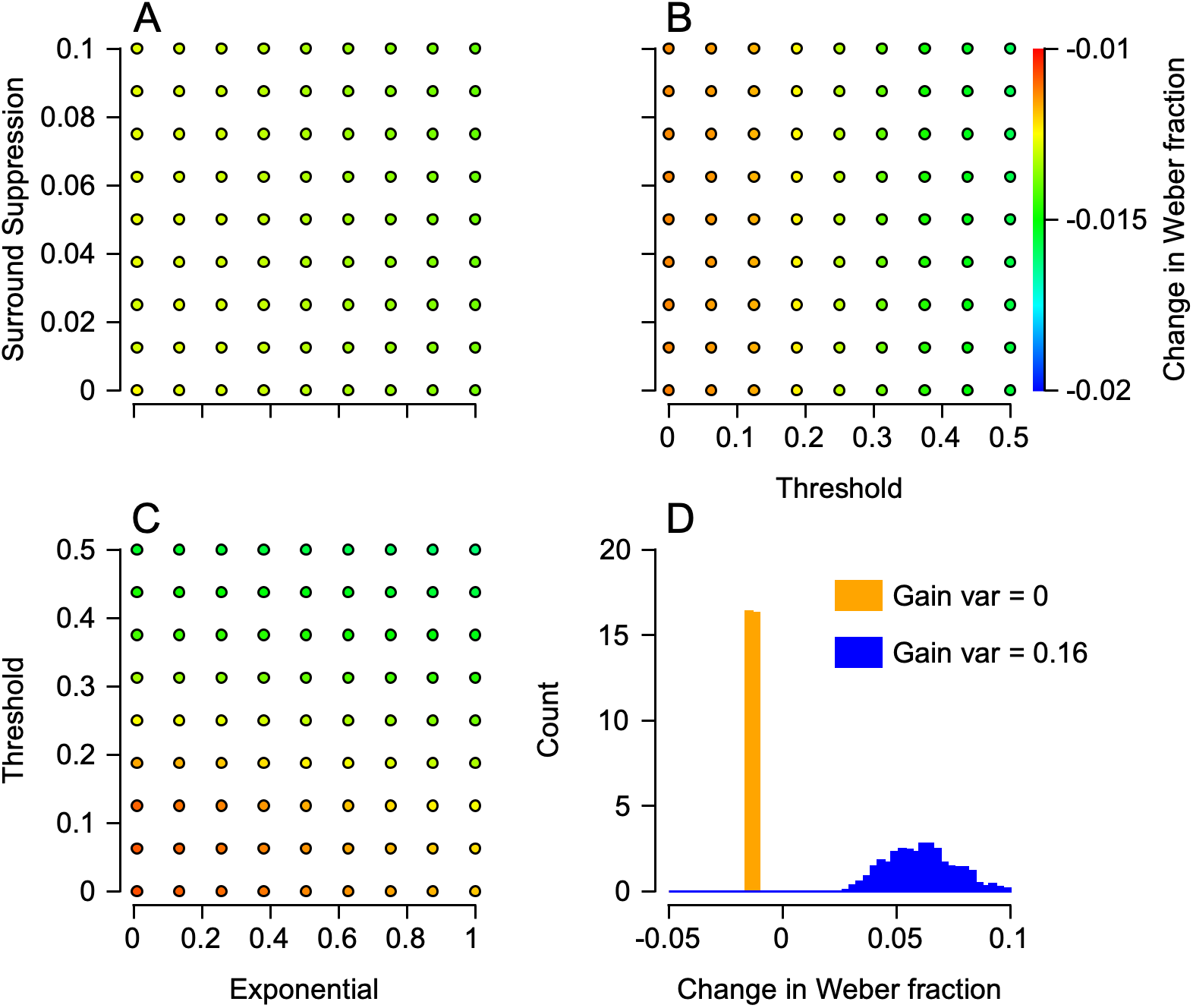
Gain noise is required to match behavior and physiology across a range of circuit model parameterizations. To determine the degree to which the parameters of MT model neurons can replicate behavioral results without gain noise, we measured circuit model behavior while systematically varying each parameter. For each model simulation, we used a specific combination of the exponential (*n*), surround suppression (*β*), and threshold (*ω*) of MT model neurons (see Methods). Across model simulations, we systematically sampled each possible parameter combination. For each simulated parameter combination, we fit a model of the form *σ*^2^ = *w*^2^ *μ*^2^ to the mean and variance of the circuit output, and allowed *w* to change with target size. Panel A plots the change in Weber fraction, measured as the difference in *w* fit to the 2 deg target and *w* fit to the 20 deg target (colors; see color bar in panel B), for each combination of exponential and surround suppression, averaged across thresholds. Panel B plots the changes in Weber fraction, as in panel A, but as a function of the threshold and surround suppression, averaged across exponentials. Panel C plots the change in Weber fraction, as in panel A, but as a function of the exponential and threshold, averaged across surround suppression. While the parameters had a small effect on the change in Weber fraction (note the narrow range in the color bar), all combinations resulted in a slightly larger Weber fraction for the 20 vs. 2 deg target, as indicated by the negative change in Weber fraction. Panel D summarizes the results by plotting the relative frequency of the measured change in Weber fractions across all parameter combinations for simulations with (blue) and without (orange) noise in the motion reliability pathway. Without noise, the Weber fractions modestly increased from the 2 deg target to the 20 deg target, as indicated by the negative values, which was inconsistent with our behavioral data. With noise, Weber fractions decreased with increasing target size, as indicated by the positive values, which was consistent with our behavioral data. Overall, the results demonstrate that no combination of model MT neuron parameters allowed circuit behavior to match the behavioral results without the addition of gain noise.

**Supplementary Figure 4.**
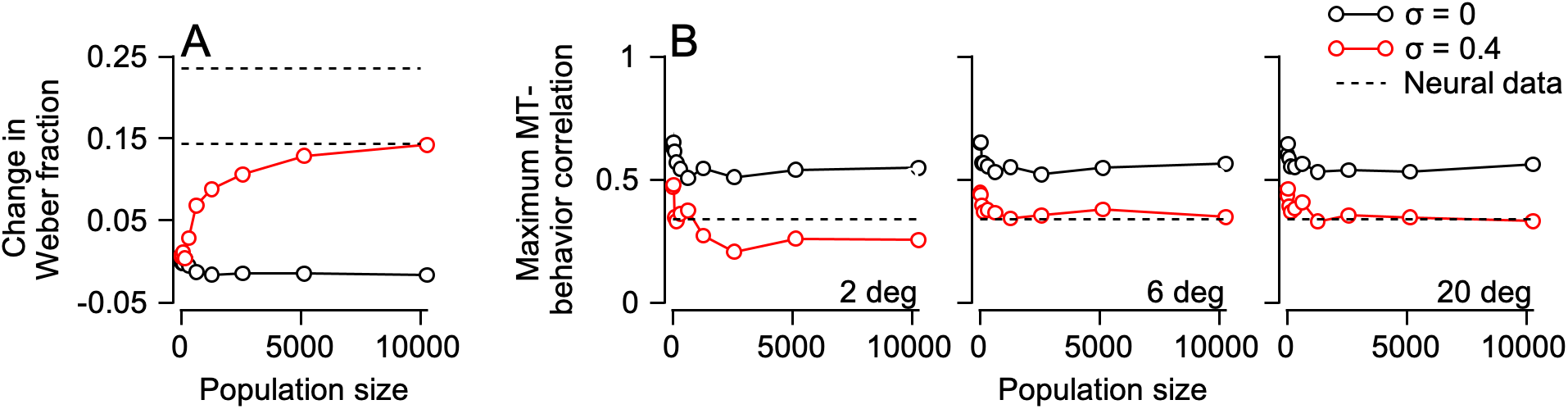
Effect of population size on circuit model results. To determine how population size impacts decoding results, we simulated circuit output with model MT neuron populations sizes of 20, 40, 80, 160, 320, 640, 1280, 2560, 5120, or 10240 with (red) or without (black) gain noise. Panel A plots the change in Weber fraction, measured as in Supplementary Figure 3, for each simulation. Only for the smallest population sizes did gain noise fail to produce a change in Weber fraction, and a change was never observed for models without gain noise. Intriguingly, only for the largest simulated population sizes did the model approach the change in Weber fraction observed in monkey behavior (dashed lines indicate the range of observed changes across monkeys and target directions). We then measured the magnitude of model MT-behavior correlations from the maximum observed MT-correlation across target speeds and model neurons. Panel B plots the magnitude for each population size for the 2 (left), 6 (middle), and 20 (right) deg targets. The magnitude of MT-behavior correlations measured from the circuit model were consistently larger than those observed in physiology (horizontal dashed line) when simulated without gain noise (black), regardless of the population size. Addition of gain noise decreased MT-behavior correlations to realistic levels for all population sizes tested (red).

## Notes

### Competing Interest Statement

The authors have declared no competing interest.

## References

1. Fitts, P. M. The information capacity of the human motor system in controlling the amplitude of movement. J. Exp. Psychol. 47, 381–391 (1954).

2. Fechner, G. T. Elements of Psychophysics, vol. 1 of Henry Holt Additions in Psychology (Holt, Rinehart and Winston, Inc., United States of America, 1966).

3. Britten, K. H., Shadlen, M. N., Newsome, W. T. & Movshon, J. A. The analysis of visual motion: a comparison of neuronal and psychophysical performance. J. Neurosci. 12, 4745–4765 (1992).

4. Priebe, N. J. & Lisberger, S. G. Estimating target speed from the population response in visual area MT. J. Neurosci. 24, 1907–1916 (2004).

5. Jazayeri, M. & Movshon, J. A. A new perceptual illusion reveals mechanisms of sensory decoding. Nature 446, 912–915 (2007).

6. Beck, J. M. et al. Probabilistic population codes for bayesian decision making. Neuron 60, 1142–1152 (2008).

7. Park, I. M., Meister, M. L. R., Huk, A. C. & Pillow, J. W. Encoding and decoding in parietal cortex during sensorimotor decision-making. Nat. Neurosci. 17, 1395–1403 (2014).

8. Funamizu, A., Kuhn, B. & Doya, K. Neural substrate of dynamic bayesian inference in the cerebral cortex. Nat. Neurosci. 19, 1682–1689 (2016).

9. Jazayeri, M. & Shadlen, M. N. Temporal context calibrates interval timing. Nat. Neurosci. 13, 1020–1026 (2010).

10. Pardo-Vazquez, J. L. et al. The mechanistic foundation of weber’s law. Nat. Neurosci. 22, 1493–1502 (2019).

11. Harris, C. M. & Wolpert, D. M. Signal-dependent noise determines motor planning. Nature 394, 780–784 (1998).

12. Todorov, E. & Jordan, M. I. Optimal feedback control as a theory of motor coordination. Nat. Neurosci. 5, 1226–1235 (2002).

13. Churchland, M. M. et al. Stimulus onset quenches neural variability: a widespread cortical phenomenon. Nat. Neurosci. 13, 369–378 (2010).

14. Churchland, M. M., Afshar, A. & Shenoy, K. V. A central source of movement variability. Neuron 52, 1085–1096 (2006).

15. Schoppik, D., Nagel, K. I. & Lisberger, S. G. Cortical mechanisms of smooth eye movements revealed by dynamic covariations of neural and behavioral responses. Neuron 58, 248–260 (2008).

16. Sober, S. J. & Sabes, P. N. Flexible strategies for sensory integration during motor planning. Nat. Neurosci. 8, 490–497 (2005).

17. Acerbi, L., Vijayakumar, S. & Wolpert, D. M. On the origins of suboptimality in human probabilistic inference. PLoS Comput. Biol. 10, e1003661 (2014).

18. Drugowitsch, J., Wyart, V., Devauchelle, A.-D. & Koechlin, E. Computational precision of mental inference as critical source of human choice suboptimality. Neuron 92, 1398–1411 (2016).

19. Newsome, W. T., Wurtz, R. H., Dürsteler, M. R. & Mikami, A. Deficits in visual motion processing following ibotenic acid lesions of the middle temporal visual area of the macaque monkey. J. Neurosci. 5, 825–840 (1985).

20. Groh, J. M., Born, R. T. & Newsome, W. T. How is a sensory map read out? effects of microstimulation in visual area MT on saccades and smooth pursuit eye movements. J. Neurosci. 17, 4312–4330 (1997).

21. Born, R. T., Groh, J. M., Zhao, R. & Lukasewycz, S. J. Segregation of object and background motion in visual area MT: effects of microstimulation on eye movements. Neuron 26, 725–734 (2000).

22. Osborne, L. C., Hohl, S. S., Bialek, W. & Lisberger, S. G. Time course of precision in smooth-pursuit eye movements of monkeys. J. Neurosci. 27, 2987–2998 (2007).

23. Nover, H., Anderson, C. H. & DeAngelis, G. C. A logarithmic, scale-invariant representation of speed in macaque middle temporal area accounts for speed discrimination performance. J. Neurosci. 25, 10049–10060 (2005).

24. Hohl, S. S., Chaisanguanthum, K. S. & Lisberger, S. G. Sensory population decoding for visually guided movements. Neuron 79, 167–179 (2013).

25. Lisberger, S. G. Visual guidance of smooth pursuit eye movements. Annu. Rev Vis Sci 1, 447–468 (2015).

26. Tanaka, M. & Lisberger, S. G. Regulation of the gain of visually guided smooth-pursuit eye movements by frontal cortex. Nature 409, 191–194 (2001).

27. Schwartz, J. D. & Lisberger, S. G. Initial tracking conditions modulate the gain of visuo-motor transmission for smooth pursuit eye movements in monkeys. Vis. Neurosci. 11, 411–424 (1994).

28. Yang, J., Lee, J. & Lisberger, S. G. The interaction of bayesian priors and sensory data and its neural circuit implementation in visually guided movement. J. Neurosci. 32, 17632–17645 (2012).

29. Darlington, T. R., Tokiyama, S. & Lisberger, S. G. Control of the strength of visual-motor transmission as the mechanism of rapid adaptation of priors for bayesian inference in smooth pursuit eye movements. J. Neurophysiol. 118, 1173–1189 (2017).

30. Darlington, T. R., Beck, J. M. & Lisberger, S. G. Neural implementation of bayesian inference in a sensorimotor behavior. Nat. Neurosci. 21, 1442–1451 (2018).

31. Lee, J., Joshua, M., Medina, J. F. & Lisberger, S. G. Signal, noise, and variation in neural and sensory-motor latency. Neuron 90, 165–176 (2016).

32. van Beers, R. J., Sittig, A. C. & Gon, J. J. Integration of proprioceptive and visual position-information: An experimentally supported model. J. Neurophysiol. 81, 1355–1364 (1999).

33. Körding, K. P. & Wolpert, D. M. Bayesian integration in sensorimotor learning. Nature 427, 244–247 (2004).

34. Liu, D. & Todorov, E. Evidence for the flexible sensorimotor strategies predicted by optimal feedback control. J. Neurosci. 27, 9354–9368 (2007).

35. Weiler, J., Gribble, P. L. & Pruszynski, J. A. Spinal stretch reflexes support efficient hand control. Nat. Neurosci. 22, 529–533 (2019).

36. Pola, J. & Wyatt, H. J. Active and passive smooth eye movements: effects of stimulus size and location. Vis. Res. 25, 1063–1076 (1985).

37. Lisberger, S. G. & Westbrook, L. E. Properties of visual inputs that initiate horizontal smooth pursuit eye movements in monkeys. J. Neurosci. 5, 1662–1673 (1985).

38. Heinen, S. J. & Watamaniuk, S. N. Spatial integration in human smooth pursuit. Vis. Res. 38, 3785–3794 (1998).

39. Osborne, L. C., Bialek, W. & Lisberger, S. G. Time course of information about motion direction in visual area MT of macaque monkeys. J. Neurosci. 24, 3210–3222 (2004).

40. Born, R. T. & Tootell, R. B. Segregation of global and local motion processing in primate middle temporal visual area. Nature 357, 497–499 (1992).

41. Liu, L. D., Haefner, R. M. & Pack, C. C. A neural basis for the spatial suppression of visual motion perception. Elife 5 (2016).

42. Zohary, E., Shadlen, M. N. & Newsome, W. T. Correlated neuronal discharge rate and its implications for psychophysical performance. Nature 370, 140–143 (1994).

43. Huang, X. & Lisberger, S. G. Noise correlations in cortical area MT and their potential impact on trial-by-trial variation in the direction and speed of smooth-pursuit eye movements. J. Neurophysiol. 101, 3012–3030 (2009).

44. de Bruyn, B. & Orban, G. A. Human velocity and direction discrimination measured with random dot patterns. Vis. Res. 28, 1323–1335 (1988).

45. Verghese, P. & Stone, L. S. Combining speed information across space. Vis. Res. 35, 2811–2823 (1995).

46. Lisberger, S. G. Visual guidance of smooth-pursuit eye movements: sensation, action, and what happens in between. Neuron 66, 477–491 (2010).

47. Born, R. T. Center-surround interactions in the middle temporal visual area of the owl monkey. J. Neurophysiol. 84, 2658–2669 (2000).

48. Lisberger, S. G. & Ferrera, V. P. Vector averaging for smooth pursuit eye movements initiated by two moving targets in monkeys. J. Neurosci. 17, 7490–7502 (1997).

49. Priebe, N. J., Churchland, M. M. & Lisberger, S. G. Reconstruction of target speed for the guidance of pursuit eye movements. J. Neurosci. 21, 3196–3206 (2001).

50. Bakhtiari, S. & Pack, C. C. Functional specialization in the middle temporal area for smooth pursuit initiation. MNI Open Res 2, 6 (2018).

51. Shadlen, M. N., Britten, K. H., Newsome, W. T. & Movshon, J. A. A computational analysis of the relationship between neuronal and behavioral responses to visual motion. J. Neurosci. 16, 1486–1510 (1996).

52. van Beers, R. J., Haggard, P. & Wolpert, D. M. The role of execution noise in movement variability. J. Neurophysiol. 91, 1050–1063 (2004).

53. Maunsell, J. H. & Van Essen, D. C. Functional properties of neurons in middle temporal visual area of the macaque monkey. i. selectivity for stimulus direction, speed, and orientation. J. Neurophysiol. 49, 1127–1147 (1983).

54. Albright, T. D. Direction and orientation selectivity of neurons in visual area MT of the macaque. J. Neurophysiol. 52, 1106–1130 (1984).

55. Van Essen, D. C., Maunsell, J. H. & Bixby, J. L. The middle temporal visual area in the macaque: myeloarchitecture, connections, functional properties and topographic organization. J. Comp. Neurol. 199, 293–326 (1981).

56. Erickson, R. G., Dow, B. M. & Snyder, A. Z. Representation of the fovea in the superior temporal sulcus of the macaque monkey. Exp. Brain Res. 78, 90–112 (1989).

57. Ferrera, V. P. & Lisberger, S. G. Neuronal responses in visual areas MT and MST during smooth pursuit target selection. J. Neurophysiol. 78, 1433–1446 (1997).

58. Tsui, J. M. G. & Pack, C. C. Contrast sensitivity of MT receptive field centers and surrounds. J. Neurophysiol. 106, 1888–1900 (2011).

59. Liu, L. D., Miller, K. D. & Pack, C. C. A unifying motif for spatial and directional surround suppression. J. Neurosci. 38, 989–999 (2017).

60. Behling, S. & Lisberger, S. G. Different mechanisms for modulation of the initiation and steady-state of smooth pursuit eye movements. J. Neurophysiol. 123, 1265–1276 (2020).

61. Krekelberg, B., van Wezel, R. J. A. & Albright, T. D. Interactions between speed and contrast tuning in the middle temporal area: implications for the neural code for speed. J. Neurosci. 26, 8988–8998 (2006).

62. MacAvoy, M. G., Gottlieb, J. P. & Bruce, C. J. Smooth-pursuit eye movement representation in the primate frontal eye field. Cereb. Cortex 1, 95–102 (1991).

63. Tanaka, M. & Fukushima, K. Neuronal responses related to smooth pursuit eye movements in the periarcuate cortical area of monkeys. J. Neurophysiol. 80, 28–47 (1998).

64. Tanaka, M. & Lisberger, S. G. Role of arcuate frontal cortex of monkeys in smooth pursuit eye movements. i. basic response properties to retinal image motion and position. J. Neurophysiol. 87, 2684–2699 (2002).

65. Shi, D., Friedman, H. R. & Bruce, C. J. Deficits in smooth-pursuit eye movements after muscimol inactivation within the primate’s frontal eye field. J. Neurophysiol. 80, 458–464 (1998).

66. Nuding, U. et al. TMS evidence for smooth pursuit gain control by the frontal eye fields. Cereb. Cortex 19, 1144–1150 (2008).

67. Osborne, L. C., Lisberger, S. G. & Bialek, W. A sensory source for motor variation. Nature 437, 412–416 (2005).

68. Brunton, B. W., Botvinick, M. M. & Brody, C. D. Rats and humans can optimally accumulate evidence for decision-making. Science 340, 95–98 (2013).

69. Churchland, A. K. et al. Variance as a signature of neural computations during decision making. Neuron 69, 818–831 (2011).

70. Musall, S., Kaufman, M. T., Juavinett, A. L., Gluf, S. & Churchland, A. K. Single-trial neural dynamics are dominated by richly varied movements. Nat. Neurosci. 22, 1677–1686 (2019).

71. Remington, E. D., Parks, T. V. & Jazayeri, M. Late bayesian inference in mental transformations. Nat. Commun. 9, 4419 (2018).

72. Egger, S. W. & Jazayeri, M. A nonlinear updating algorithm captures suboptimal inference in the presence of signal-dependent noise. Sci. Rep. 8, 12597 (2018).

73. Xiao, D. K., Raiguel, S., Marcar, V. & Orban, G. A. The spatial distribution of the antagonistic surround of MT/V5 neurons. Cereb. Cortex 7, 662–677 (1997).

74. Mukherjee, T., Liu, B., Simoncini, C. & Osborne, L. C. The spatiotemporal filter for visual motion integration from pursuit eye movements in humans and monkeys. J. Neurosci. 37, 1394–1412 (2016).

75. Landy, M. S., Maloney, L. T., Johnston, E. B. & Young, M. Measurement and modeling of depth cue combination - in defense of weak fusion. Vis. Res. 35, 389–412 (1995).

76. Ernst, M. O. & Banks, M. S. Humans integrate visual and haptic information in a statistically optimal fashion. Nature 415, 429–433 (2002).

77. Battaglia, P. W., Jacobs, R. A. & Aslin, R. N. Bayesian integration of visual and auditory signals for spatial localization. J. Opt. Soc. Am. A Opt. Image Sci. Vis. 20, 1391–1397 (2003).

78. Oruç, İ., Maloney, L. T. & Landy, M. S. Weighted linear cue combination with possibly correlated error. Vis. Res. 43, 2451–2468 (2003).

79. Alais, D. & Burr, D. The ventriloquist effect results from near-optimal bimodal integration. Curr. Biol. 14, 257–262 (2004).

80. Mamassian, P. & Landy, M. S. Observer biases in the 3D interpretation of line drawings. Vis. Res. 38, 2817–2832 (1998).

81. Weiss, Y., Simoncelli, E. P. & Adelson, E. H. Motion illusions as optimal percepts. Nat. Neurosci. 5, 598–604 (2002).

82. Todorov, E. Optimality principles in sensorimotor control. Nat. Neurosci. 7, 907–915 (2004).

83. Scott, S. H. Optimal feedback control and the neural basis of volitional motor control. Nat. Rev. Neurosci. 5, 532–546 (2004).

84. Ramachandran, R. & Lisberger, S. G. Normal performance and expression of learning in the vestibulo-ocular reflex (VOR) at high frequencies. J. Neurophysiol. 93, 2028–2038 (2005).

85. Britten, K. H. & Heuer, H. W. Spatial summation in the receptive fields of MT neurons. J. Neurosci. 19, 5074–5084 (1999).

86. Rust, N. C., Mante, V., Simoncelli, E. P. & Movshon, J. A. How MT cells analyze the motion of visual patterns. Nat. Neurosci. 9, 1421–1431 (2006).

87. Pack, C. C., Hunter, J. N. & Born, R. T. Contrast dependence of suppressive influences in cortical area MT of alert macaque. J. Neurophysiol. 93, 1809–1815 (2005).

88. Cicchini, G. M., Arrighi, R., Cecchetti, L., Giusti, M. & Burr, D. C. Optimal encoding of interval timing in expert percussionists. J. Neurosci. 32, 1056–1060 (2012).

